# A millennium of increasing ecosystem diversity until the mid-20^th^ century

**DOI:** 10.1101/2021.03.16.435590

**Authors:** Inês S. Martins, Maria Dornelas, Mark Vellend, Chris D. Thomas

## Abstract

Land-use change is widely regarded as a simplifying and homogenising force in nature. In contrast, analysing global land-use reconstructions from the 10^th^ to 20^th^ centuries, we found progressive increases in the number, evenness, and diversity of ecosystems (including human-modified land-use types) across the globe. Ecosystem diversity increased more rapidly after ∼1700CE, then slowed or partially reversed (depending on the metric) following the mid-20^th^ century acceleration of human impacts. Differentiation also generally increased across space, with homogenization only evident in the presence-absence analysis of ecosystem types at the global scale. Our results suggest that human land-use changes have primarily driven increases in ecosystem diversity over the last millennium.

## Main Text

Humans have been reshaping the processes, structure and biological composition of ecosystems for millennia. These changes are typically regarded as the most important proximate drivers of terrestrial biodiversity change: the International Union for Conservation of Nature (IUCN) lists various aspects of land-use change and altered management as eight of the top ten threats to species (*1*), while the Intergovernmental Science-Policy Platform on Biodiversity and Ecosystem Services (IPBES) identifies ‘changes in land and sea use’ as the largest driver of ‘changes in nature’ (*2*). These changes are recognised by the United Nations Convention on Biological Diversity (CBD), which has ecosystem extent at the core of its post-2020 agenda (*3*).

Ecosystem diversity is regarded both as a key component of biodiversity in its own right by the CBD, which recognises biodiversity as encompassing the “*diversity within species, between species and of ecosystems*” (*4*), and as a key determinant of species richness (*5*). Despite the central importance of ecosystem diversity changes, there is no comprehensive analysis of how land-use change has altered the diversity of ecosystem types over time and space. Almost all attention in the literature has been on changes in the coverage of specific ecosystem types (*2*) rather than ecosystem diversity *per se*. It is unclear whether land-use change has generally led to landscape simplification (e.g. as in some extensive arable landscapes) or landscape diversification (a greater mixture of ecosystems). Ecosystem diversity and its changes over time represent a major gap in our broader understanding of human impacts on biodiversity.

Ecosystems are difficult to define unambiguously, so we here designate land cover types that contain distinct plant-based physical structures and their associated biotas as “ecosystems”. For example, primary forest (natural), rangelands (semi-natural) and arable (anthropogenic) land covers are all included within this definition; and we regard a landscape that contains all three as having greater ecosystem diversity than those that only contain one of them (see Methods). We used global reconstructions of land-use data from the LUH2 dataset (*6*) to estimate the coverage of 9 different anthropogenic and relatively natural ecosystems at 0.25° resolution (∼769km^2^, or ∼27.8km x 27.6km ‘landscapes’ at the equator), as used recently as inputs to the modelling of past and future biodiversity change (*7*). These data layers are built on multiple model inputs (e.g. using existing local/regional level land statistics or records, population and cultural reconstructions, historical maps), with uncertainty expected to be highest in the distant past. For this reason, we assess changes in ecosystem diversity in time steps of 100 years since 900CE and decade-long time steps from 1700 to 2000, when data quality improved (see Methods).

As expected, the data show declining areas of primary forested and primary non-forested land, a higher rate of decline after 1700, and growing areas of multiple human-dominated land uses (Fig. S1). Surprisingly, the *frequencies* of areas that include primary forested and primary non-forested land remain largely unaltered. That is, most (96.6% for forests and 96.0% for non-forested land) of the 0.25° cells where these ecosystem types were present in the 10^th^ century still contained at least some area of the same ecosystem type in the 20^th^ century (Fig. S1; Data S1). This means that while we have seen substantial post-1700 declines in areas of primary land cover, these ecosystem types still persist almost everywhere.

We quantified changes to the diversity of ecosystems at different scales using six complementary metrics. The first four are metrics of ecosystem diversity changes through time within 0.25° cells (*α* diversity) and the remaining two are comparisons across space as well as time, measuring whether the ecosystem composition in different 0.25° cells is becoming more or less similar over time (*β* diversity). The metrics we use are equivalent to those typically applied to measure species diversity, but with different ecosystems in place of different species, and the area of an ecosystem equivalent to the abundance of a given species.

The four ecosystem *α* diversity metrics were: ecosystem richness (number of ecosystem types present per cell), Pielou’s evenness Index (the balance among relative areas of each ecosystem type), the Shannon diversity Index (heterogeneity, combining richness and evenness) and Rao’s quadratic entropy Index (incorporating the richness, evenness, and biological distinctiveness of each ecosystem type, for which reason it is the metric likely to correlate most strongly with species diversity) (see Methods). All four indices showed significant (based on bootstrapping) and continuous increases since the 10^th^ century, with a clear increase in rates from 1700 onwards, evident on both the centurial and decadal time scales (Fig. 1). Mean global ecosystem richness increases are predominantly driven by a reduction in the number of cells containing one to four ecosystem types, and increases in areas supporting five or more ecosystem types (Fig. 1A inset). This is a consequence of the frequencies of different anthropogenic ecosystems (e.g. pastures, cropland) growing considerably faster than the frequencies of primary vegetation decline (Fig. S1). The net effect of modification has been to increase ecosystem diversity: an increased number, evenness, heterogeneity and compositional entropy of ecosystem types.

**Fig. 1.**
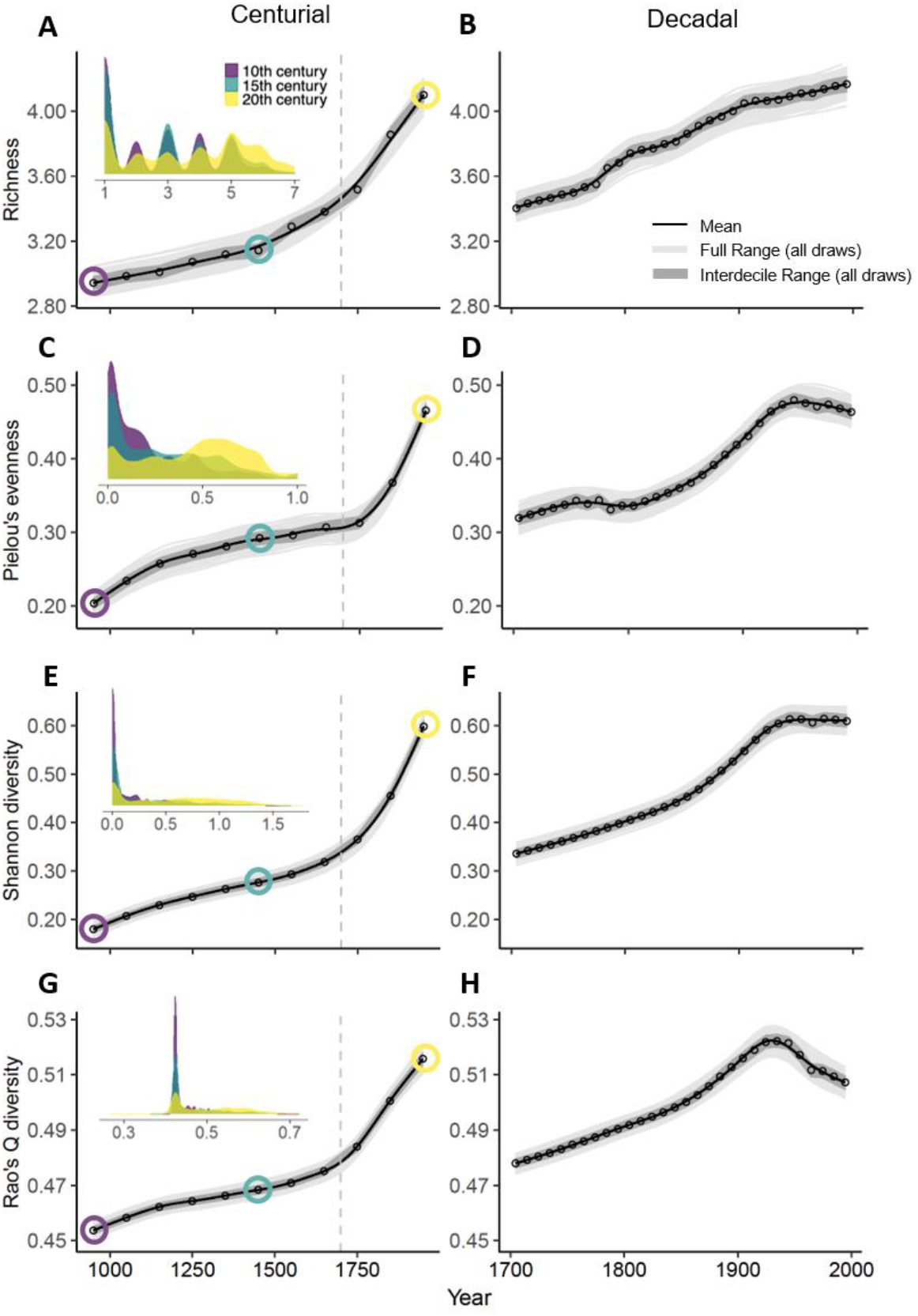
Net change in local ecosystem diversity from 900 to 2000. **(A-B)** within-cell ecosystem richness (mean numbers of ecosystem types per cell), **(C-D)** Pielou’s evenness (J) **(E-F)** Shannon diversity Index (H’) **(G-H)** and Rao’s quadratic entropy Index (RaoQ). C-D and G-H scale from 0 (minimum) to 1 (maximum evenness and diversity). The left-hand graphs show the centurial trends (from 900 to 2000), where each point on the graph represents the spatial averaged means across all cells (0.25° x 0.25°) found over a 100 year period plotted on the mid-point of the century, while the right-hand graphs show the decadal averages (from 1700 to 2000) plotted on the mid-point of the decade. The smoothed lines represent fitted GAMs (generalized additive models), the dark grey shading represents the interdecile range calculated from 1000 sampling draws (each draw contained 2095 random sampling cells -full range of draws is shown in light grey). Inset density plots on the left show the distribution of the individual cell estimates of the different metrics at 3 points in time.

This trend of increasing diversity at the 0.25° cell resolution changes after the mid-20^th^ century: the rate of increase slows for ecosystem richness, flattens for Shannon heterogeneity, shows a possible downturn (but shallower than the interdecile range) for evenness and reverses for Rao’s quadratic entropy Index (Fig. 1). This is coincident with the ‘Great Acceleration’ of the human population, technologies and associated impacts (*8, 9*). Rao’s Index was the only metric to decline significantly (Fig. 1H), reflecting the reduced biological distinctiveness of different anthropogenic ecosystems types (i.e. they often share species with one another)(*10*), whose cover increased during this period (Fig. S1). However, this result was scale-dependent: Rao’s Index only reversed at sub-regional scales (< 4° cell resolution, < ∼197,000km^2^ at the equator -Figs. S2-S3). Larger areas continued to accumulate increased ecosystem diversity for all four metrics, but at a reduced rate (Fig. S3). Despite some downturns, average ecosystem diversity for the 20^th^ century remained higher than the averages of any preceding century for all metrics at all spatial scales considered (0.25° to 15° cells).

This pattern of increasing ecosystem diversity holds qualitatively for different regions of the world (IPBES sub-regions and WWF biomes), albeit with geographic variation. Rao’s quadratic entropy Index, for example, revealed net diversity increases since the 10^th^ century, rate increases from 1700 onwards, and a mixture of slow-downs and reversals in the 20^th^ century across most regions and biomes of the world (Fig. 2; see Figs. S4-S5 for other diversity metrics). Nearly all IPBES regions (16 out of 17) showed a net increase in accumulated ecosystem diversity using this metric between the 10^th^ and 20^th^ centuries, and a majority (13 out of 17) did so between 1700 and 2000, but with high interdecile variation (Figs. S4-S5). Divergent trajectories likely reflect the timing of different human impacts in different regions (e.g. Fig. S6).

**Fig. 2.**
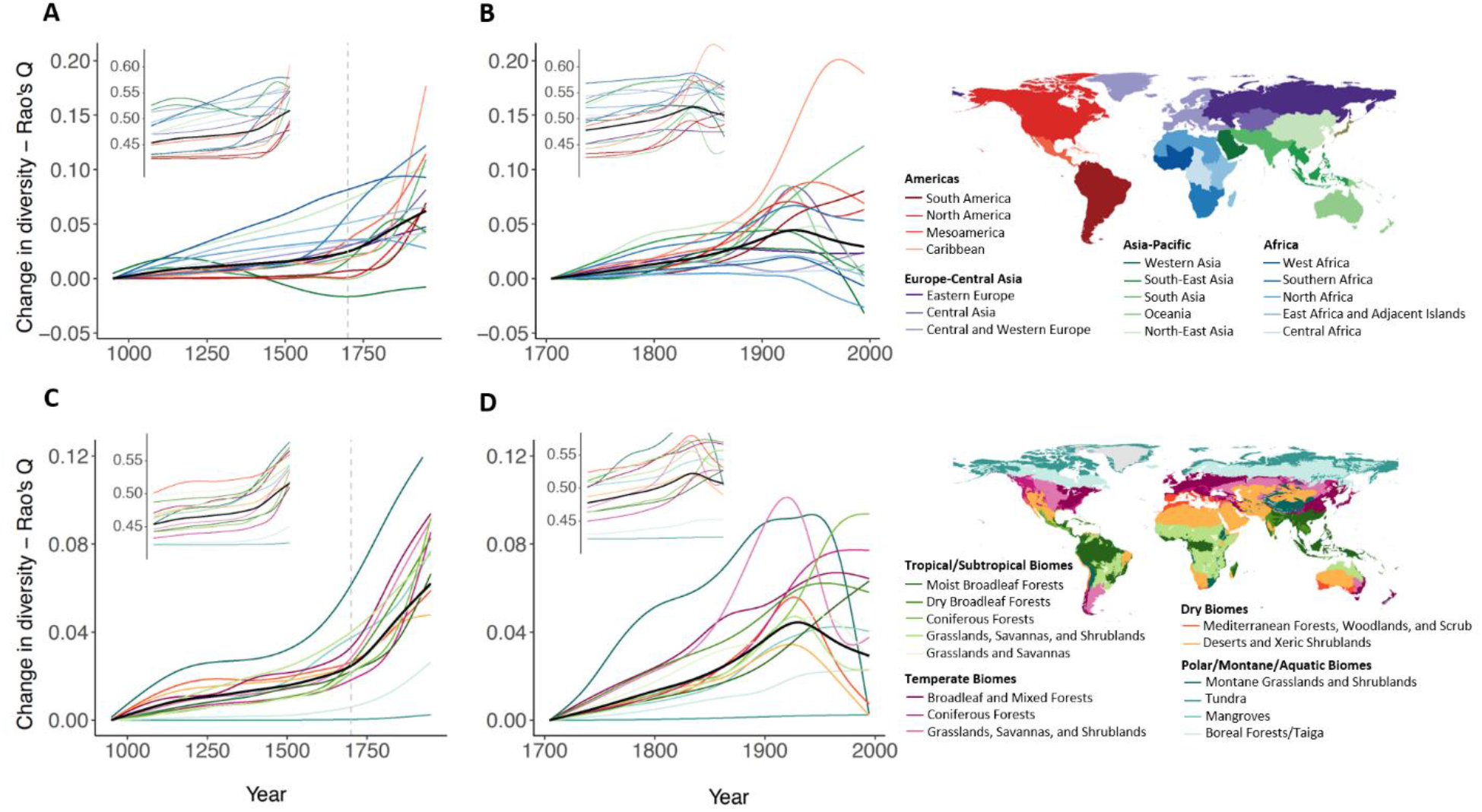
Regional changes in local ecosystem diversity (*α* diversity). Main plots show the GAM-fitted trends for **(A-B)** each IPBES sub-region and **(C-D)** each WWF Biome, measured by Rao’s Q Index. Centurial trends (A, C) are shown relative to the 10^th^ century, and decadal trends (B-D) relative to 1700. Insets show absolute values. Colours represent different regions. Black lines show GAM-fitted global trends. Other diversity metrics are shown in Figs. S4-S5.

Our two measures of spatial diversity (*β* diversity) also tended to increase over time, although the results varied with the grain (size of cells) and extent (maximum distance between pairs of cells being compared) of analysis. The Jaccard dissimilarity index, which measures differences in which ecosystem types are present (incidence) in different locations, revealed a pattern of increasing differentiation over time at sub-global scales (within ≤ 9° cells ∼997,000km^2^ at the equator, or approximately the size of Egypt) but homogenization at a global extent (Fig. 3, Figs. S7-S8). The latter is consistent with some individual ecosystem types becoming present in more cells across the globe (e.g. at least small areas of croplands are found in large numbers of cells). In contrast, the Bray-Curtis dissimilarity index, which incorporates the area of each ecosystem type as well as which ecosystem types are present, showed growing differentiation over time across all scales (grains and extents; Fig. 3, Figs. S7-S8). Increasingly, some locations have high percentages and others low percentages of particular ecosystem types. There was a possible slight shift in rates of change of spatial differentiation in the mid-20^th^ century, but no reversals of previous trends (Fig. S8).

**Fig. 3.**
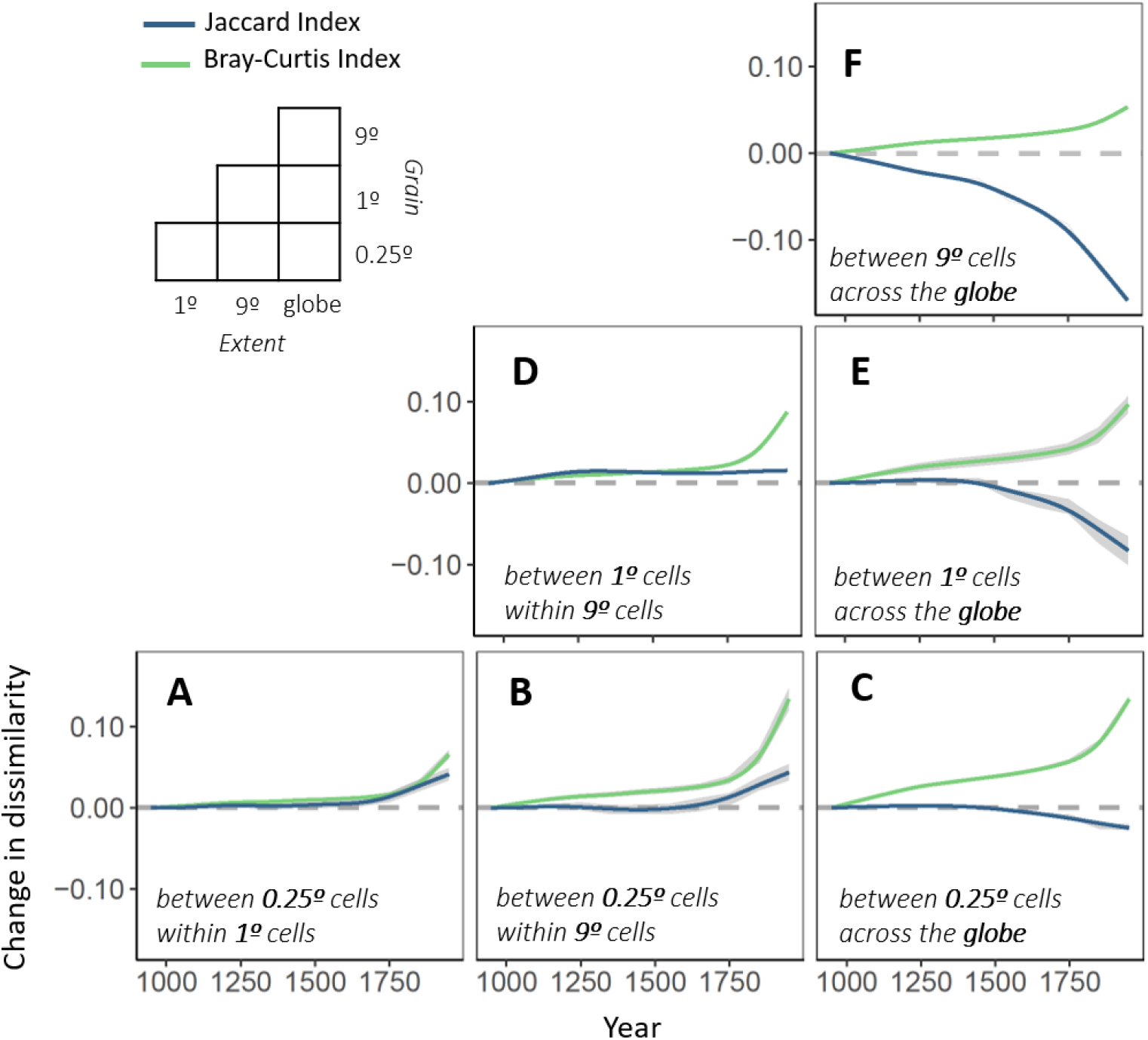
Temporal trends in spatial *β* diversity of ecosystems. (**A-F**) Average total dissimilarity change between pairs of smaller cells (grain) within increasing larger cell areas (extent) as measured by the Jaccard Index (ecosystem type presence-absence dissimilarity -blue lines) and Bray-Curtis Index (ecosystem type presence-absence and area coverage dissimilarity -green lines) between the 10^th^ century and each subsequent century. Upper left-side legend shows the different grains of analysis and extents considered. The smoothed lines represent fitted GAMs (generalized additive models). For (A, B, C, E) diversity change is characterized by the average dissimilarity change from 100 random draws (each draw ∼1% of the full dataset), with the interdecile range calculated from all draws (dark grey shading). The decomposition of total dissimilarity into its components is shown in Fig. S7 (centurial) and Fig. S8 (decadal).

The changing diversity and distributions of ecosystems represent a major global change, of importance to regional and global-scale ecosystem processes, to functions, and to the provision of services (*2*). These changes underlie the CBD aspiration to achieve “*no net loss”* of ecosystems by 2030 (*3*). However, ‘lost’ ecosystems have been replaced by a variety of anthropogenic ecosystems, increasing the numbers of ecosystem types per 0.25° landscape in most parts of the world (Figs. 1-2), and also increasing spatial differentiation within most country-sized regions (Fig. 3). Many of these transformations are of great antiquity, reflecting the diversity of the peoples who inhabited them, making these parts of the planet less hospitable to some species but more so for others. In fact, ongoing conservation programmes commonly highlight the human and biodiversity value of cultural and indigenous landscapes in all six populated continents (*11*–*13*). Articulating all of these ecosystem changes as ‘loss’ does not capture the full range of realities of the transformed Anthropocene world, and should be replaced by a narrative of ecosystem ‘change’.

The replacement of species-rich ecosystems by anthropogenic ecosystems (land-use cover types) that support impoverished biotas can potentially result in a loss of local (e.g. per m^2^ or per ha) species richness (*10*), and locations that today share anthropogenic ecosystem types may share increasing numbers of species (*14*). Anthropogenic ecosystems can promote the establishment of already-widespread species (*15*–*17*), while more narrowly distributed native species decline (*18*). Such conclusions have led to an overall narrative of ecosystem and biodiversity ‘decline and homogenisation’. However, the increasing diversity of ecosystem types that we observe could have the opposite effect at a landscape or regional scale (e.g. in 0.25° cells), given that ecosystem diversity is a major determinant of total species richness (*5*). After land-use change, many native species survive in the remaining areas (fragments in some places) of original ecosystems, while additional colonising species establish in different semi-natural and anthropogenic ecosystems (*19, 20*). This contrast between plot-scale species-richness results (*10*) and our landscape-scale ecosystem diversity results may help explain why observed biodiversity changes are scale-dependent (*14, 21*), and include locations and periods with increases as well as declines in numbers of species (*14, 22*–*27*). These contrasting trends come together within the Rao Index results (Figs. 1-2), which incorporates the diversity of ecosystems but ‘down-weights’ ecosystem types that often support overlapping biotas (e.g. cropland and pasture). Rao’s Index shows strongly increasing diversity over the full period, but it is also the one metric to exhibit a clear decline in diversity in the second half of the 20^th^ century.

In conclusion, we observe net ecosystem diversification across the globe over the last millennium of human transformations of the Earth’s ecosystems, and spatial differentiation at sub-global scales -prior to the mid-20^th^ century. While the global story of biodiversity change involves the loss and decline of certain ecosystem types, the full story is more complex and interesting, involving gains and increases in other ecosystem types, and increased ecosystem diversity at most spatial and temporal scales. It therefore seems appropriate to replace language emphasizing only habitat and ecosystem ‘loss’ with descriptions of ecosystem ‘change’.

## Methods

### Land-use data

We consider the last millennium, given the antiquity of many land-use changes, and we explore changes at a global scale to avoid the risk of selecting unrepresentative regions. Of candidate datasets of sufficient duration (including (*6, 28-31*), only the newly released Land Use Harmonization version 2 (LUH2) dataset (http://luh.umd.edu/data.shtml) had sufficient spatial resolution, temporal resolution *and* land-use thematic resolution for us to be able to perform the analyses. We downloaded the LUH2 global annual gridded maps (0.25° x 0.25° cell resolution) that provide the fraction of each of 12 land-use types in each cell for historical land-use change (from 900-2000, the eleven full century-long periods within the database). The 12 land-use categories were: primary vegetation (forest and non-forest), secondary vegetation (forest and non-forest), managed pasture, rangeland, urban land, plus 5 functional crop categories (including plantations). For analysis, we grouped the crop land-use data into 2 major land-use classes, cropland and tree plantations, in line with our structural definition of ecosystem types; using the same categorization as recently used in Chapter 4 of the IPBES global assessment (*7*). Because the structure, biological composition and potential fates of ecosystems vary geographically, we carried out separate analyses for each of 17 IPBES sub-regions and each of 14 WWF biomes (and also tested for effects of spatial scale, see below), in addition to a global analysis of land-use categories. These sub-global analyses test for the robustness of the results to geographic region, and to the definition of ecosystem (i.e. land-use*region combinations effectively represent a narrower definition of ecosystem type).

Data for years within centuries up to 1700, and years within decades from 1700 to 2000, were not independent of one another (see Fig. S9 and “*Temporal trends in αdiversity” section* for more details), and hence we generated two temporal datasets: centurial means for 900 to 2000, and decadal means for 1700 to 2000.

Because of sparse spatial information for some parts of the world, the LUH2 dataset is known to have some allocation issues for primary and secondary (forest and non-forest), especially in northern Africa. This was not seen to be a significant issue for our analysis (Fig. S10), and thus cells across all regions of the world were considered. However, we removed cells on the WWF Biomes boundaries, since the precise locations of such boundaries are not static (their distribution is tightly linked to dynamic climatic and geological processes), and likely to have changed over the last millennium (*32*). The final dataset contained 210363 cells at the 0.25° cell resolution. Presence, area and coverage of each ecosystem were calculated for each 0.25° grid cell (Fig. S1) and averaged at different spatial scales (grain and extent). All statistical analyses were performed in RStudio 1.2.1335 (*33*).

### Ecosystem diversity metrics (*α* diversity)

Our use of the term ‘ecosystem diversity’ is equivalent to most uses of the terms ‘habitat diversity’, ‘habitat heterogeneity’ and ‘landscape heterogeneity’, encompassing the variety of major vegetation types (and the animals associated with then) in a specified area or region. Ecosystem diversity was assessed for each grid cell in every time step, adapting four metrics that are typically applied to the estimation of species diversity. These are: richness (the number of ecosystems per cell), evenness (balance of ecosystem types), heterogeneity (number and relative area coverage of ecosystems) and composition (number, relative area coverage and the compositional distance between ecosystems). Specifically, within-cell richness was calculated as the number of unique ecosystem types present at a given grid cell. Evenness and heterogeneity estimates were computed using the Pielou’s evenness Index (*J*) and Shannon diversity Index (*H’*), respectively. The Pielou’s evenness Index (J) measures the extent to which the area of two or more ecosystems are similar (calculable for all cells containing two or more ecosystem type), and increases with increased evenness, where *0* ≤ *J* ≤ *1*. Shannon diversity Index (H’) diversity takes into consideration both the number of ecosystems present, and the area of each, thus jointly reflecting the two major contributions to diversity (the number of ecosystem types and the area of each), increasing with increased diversity. Both indexes were computed using R (‘vegan’ package). These three metrics describe the diversity of ecosystems directly.

The number of species that can be accommodated within a region (grid cell) depends on the distinctiveness of the biota associated with each ecosystem type (i.e. dissimilarity in species composition between ecosystems), in addition to the number ecosystems present, and their evenness. Therefore, our fourth metric of within-cell ecosystem *α* diversity is Rao’s quadratic entropy Index (*RaoQ*, where *0* ≤ *RaoQ* ≤ *1*), where we incorporated all 3 aspects by considering the expected biotic dissimilarities among ecosystems (using a distance matrix for species compositional dissimilarities between ecosystem types), weighted by the area of each ecosystem type. Species dissimilarity matrices were derived from Newbold *et al*. (*10*). RaoQ estimates were computed using the function *rao*.*diversity* in R (‘SYNCSA) package.

### Temporal trends in *α* diversity

We defined time steps *t* as 100yr (for 900 to 2000) and 10yr (for 1700 to 2000) periods, reflecting the original temporal resolution of the underlying land-use data before and after 1700 (Fig. S9). These provided temporal trends in each metric (Fig. 1), as well as changes (Fig. 2) since the first time-step (*t*_*x*_ − *t*_1_) of both time series (where *t*_1_ =900 and 1700, respectively).

While the primary spatial grain for analysis was 0.25°, we also evaluated whether the change of ecosystem diversity varied with scale by conducting separate analysis using grain size cells of 1°, 4°, 9° and 15° (Fig. S2-S3). For each time period, the grid-based estimates for each metric were spatially averaged across the globe, and separately averaged for broad biogeographic regions (IPBES sub-regions and WWF biomes). Spatial averages were weighted by the land-use area of grid cells (which vary latitudinally, and because of land/water cover). All estimates can be found in Data S2.

We visualized the uncertainty in the global and regional means by drawing 1000 random samples of *n* cells from our grid dataset (*x*_1_…, *x*_*n*_), equally distributed across all IPBES sub-regions (to avoid over/under-sampling regions), and calculating for each metric the *i* sample means (*M*_1_…, *M*_*i*_) across cells, where *M* = *f*(*x*_1_…, *x*_*n*_). The means of the different samples are shown in Fig. 1-2 (light grey lines). For all draws, sample *n*was 2095 cells, which represents 1% of the total number of cells. This *n* allows us to have a rich enough sample to represent (describe) the population, but small enough to avoid significant duplication (on average, only 21 cells were shared between pairs of draws) and most spatial-autocorrelation issues. The global means of the different samples are shown in Figures 1-2 (light grey lines), whereas the regional means of the different samples are shown in Figures S4-S5 (for IPBES sub-regions and WWF biomes, respectively). The latters were calculated by averaging, for each sample, the cells that fall within each region.

### Ecosystem diversity metrics (*β* diversity)

We measured *β* diversity as the dissimilarity among pairs of cells using two indices: an incidence-based index (Jaccard dissimilarity Index) and an abundance-based index (Bray-Curtis dissimilarity Index), again extending species-level diversity metrics to ecosystems. The Jaccard index, as applied here, estimates the extent to which any pair of locations (grid cells) share ecosystem types (0 for exactly the same ecosystems present to 1 for no overlap in the ecosystem types present). We used the ‘betapart’ package in R to calculate and decompose the incidence-based dissimilarity metric (Jaccard-*β*_*jac*_) into turnover (*β*_*jtu*_) and nestedness (*β*_*jne*_) components.

The Bray-Curtis (*β*_*bc*_) dissimilarity metric takes into account differences in the cover of each ecosystem type between pairs of cells, as well as differences in the identities of each ecosystem type; decomposed into balanced variation (*β*_*bc*−*bal*_-the areas of some ecosystems decline and other ecosystems increase from one cell to another) and any abundance gradient (*β*_*bc*−*gra*_-the areas of all ecosystems decline or increase equally from one cell to the other) (*34*). Bray-Curtis values also vary from 0 (two cells are identical in which ecosystems are present and in the areas of each ecosystem) and 1 (no ecosystems in common).

### Temporal trends in spatial *β* diversity of ecosystems

We characterized spatial *β* diversity change as the average pairwise dissimilarity of ecosystem composition between pairs of grid-cells (equation 1). Average pairwise dissimilarity is known to be a robust measure of spatial heterogeneity because it estimates the expected difference between a random pair of sites (*35*). The spatial scaling of *β* diversity is also important because the metrics indicate ecosystem differences between locations. Therefore, we again varied the grain of analysis (size of the cells), and also how dissimilarity changes with average distance (extent, which we varied by sampling pairs of smaller cells (0.25°, 1°, 9°) within larger cell areas, up to global extent.

For each combination of grain and extent, we calculated century-and decadal-long spatial dissimilarity values between pairs of the smaller cells. Then, for each century/decade, global average pairwise dissimilarity was calculated by averaging all pairwise comparisons estimates:

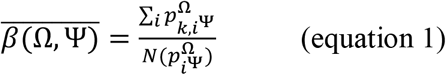

where 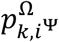 is the dissimilarity between the pair of cells ***i*** (of grain size **Ψ**) in the spatial sampling window *k* of size Ω (extent), and 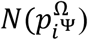 is the total number of pairwise comparisons across all windows of size Ω. Note that for each sub-global extent, pairwise dissimilarities were only calculated for pairs of smaller cells that occur within a given larger cell (e.g. between Ψ = 0.25° cell grain, within Ω= 1° x 1° extent), while at the greatest extent (Ω = global) pairwise dissimilarity was calculated for any possible pair of smaller cells across the globe. Finally, we report dissimilarity change between the first time-step (900 and 1700) and subsequent time-steps (*t*_*x*_−*t*_1_) for the two temporal scales (all estimates are presented in Data S3).

For certain combinations of grain and extent (grain of 0.25° at any extent and 1° at the global extent – Fig. 3 A-C, E) calculating pairwise dissimilarities for all possible pairs was computationally intractable. For these combinations, we randomly selected a subset of 2095 cells (∼1% of the full dataset), made pairwise comparisons 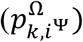 between those cells, and then estimated 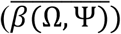. We repeated this exercise for 100 random draws, providing a mean (of the 100 draws) dissimilarity change and interdecile range (among the 100 draws). The final number of pairwise comparisons for each combination of grain and extent are shown in Data S3.

## Data and code availability

All original land-use data used is published and publicly available at https://luh.umd.edu/data.shtml. Code for all data processing and analysis and summary datasets are archived online at Zenodo (https://doi.org/10.5281/zenodo.4557542).

## Acknowledgments

We are grateful to Tim Newbold for advice on data analysis strategies, and to Anne Magurran, Jane Hill for comments on the manuscript. This work was funded by a Leverhulme Trust Research Centre -the Leverhulme Centre for Anthropocene Biodiversity. ISM also acknowledges funding from the European Union’s Horizon 2020 research and innovation program under Marie Sklodowska-Curie grant agreement no. 894644. MV was supported by the Natural Sciences and Engineering Research Council of Canada.

## Author contributions

Conceptualization: I.S.M., M.D., M.V. and C.D.T.; Methodology: I.S.M., M.D., M.V. and C.D.T.; Software: I.S.M.; Formal Analysis, I.S.M.; Visualization: I.S.M.; Writing – Original Draft: I.S.M. and C.D.T.; Writing –Review & Editing: I.S.M., M.D., M.V. and C.D.T.; Supervision: M.D., M.V. and C.D.T.; Funding Acquisition: C.D.T.

## Supplementary Information

Further information is available in the supplementary file linked to this paper. Supplementary data S1-S3 is archive online at Zenodo.

