## Supplementary Information for "A millennium of increasing ecosystem diversity until the mid-20^th^ century"

### Supplementary Information for "A millennium of increasing ecosystem diversity until the mid-20th century"

**This PDF file includes:**

Figs. S1 to S10

Captions for Data S1 to S3

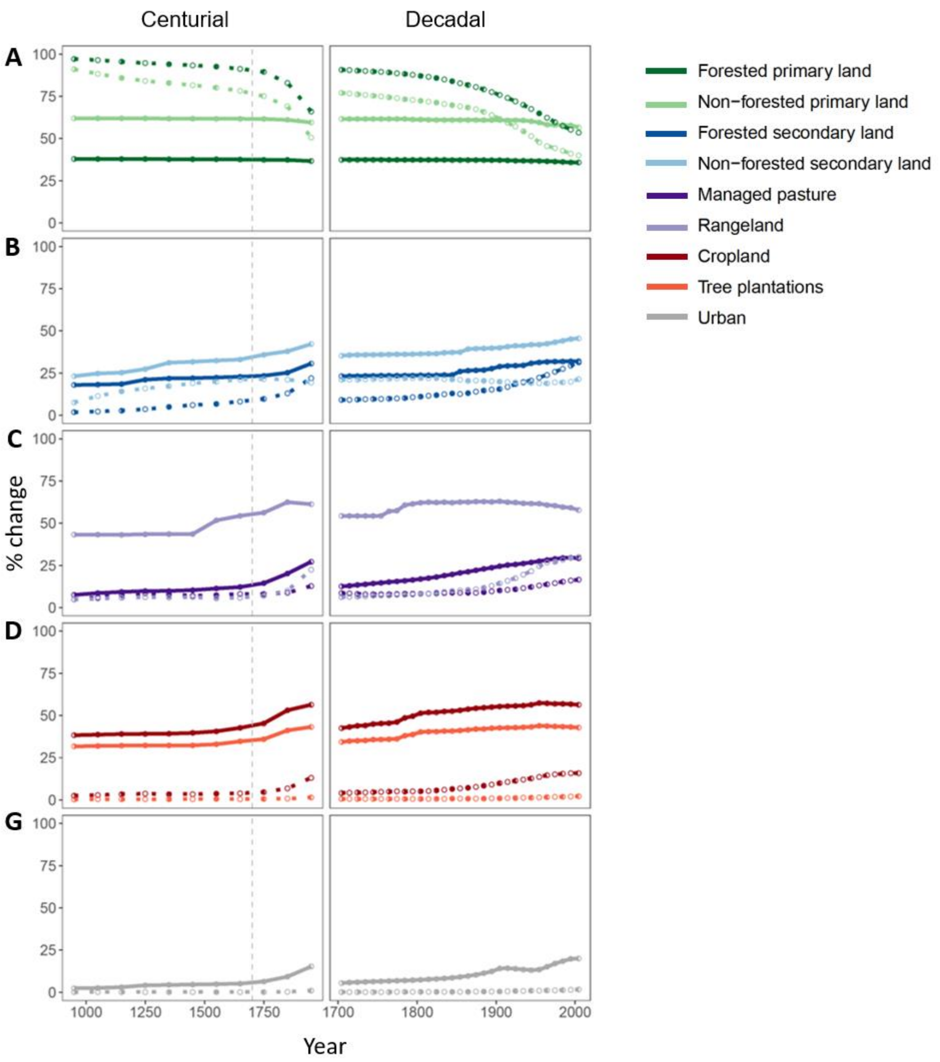


**Fig. S1. Global trends in land-use change for the nine land-use classes considered, from 900 to 2000.** Plot shows average presence (i.e. % of cells where the ecosystem is present - solid line) and coverage (i.e. % of the cell occupied by the ecosystem dashed line) of **(A)** primary land, **(B)** secondary land, **(C)** rangeland and pasture, **(D)** agricultural, and **(E)** urban ecosystems. All lines represent average values across all 0.25º x 0.25º grid cells considered (left-hand graphs show centurial means; right-hand graphs show decadal means).

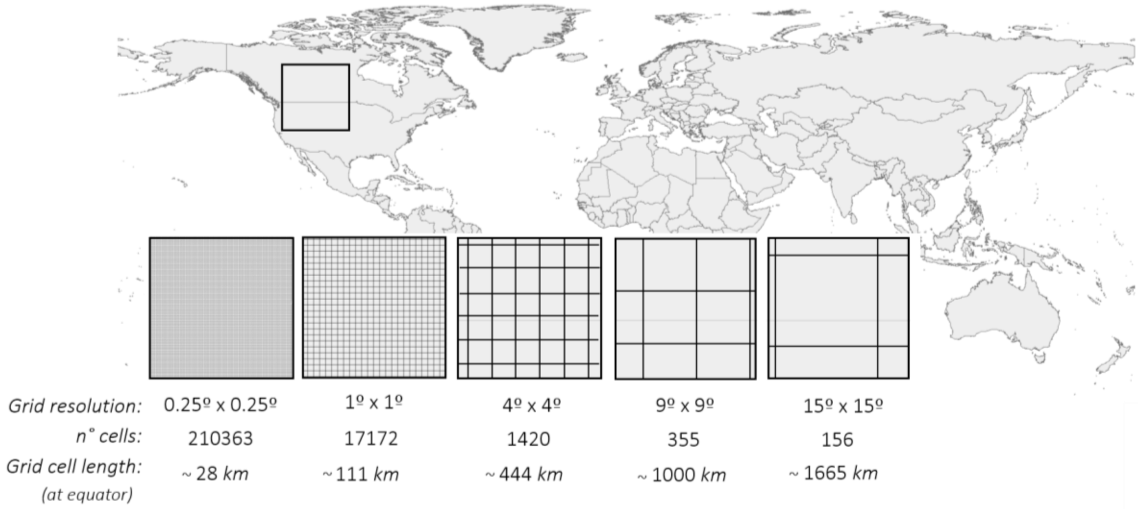


**Fig. S2. Graphical illustration of the study grains of analysis.** The primary spatial grain for analysis was 0.25º, but scale-dependence was evaluated by conducting separate analysis using grain size cells of increasing size (1º, 4º, 9º and 15º).

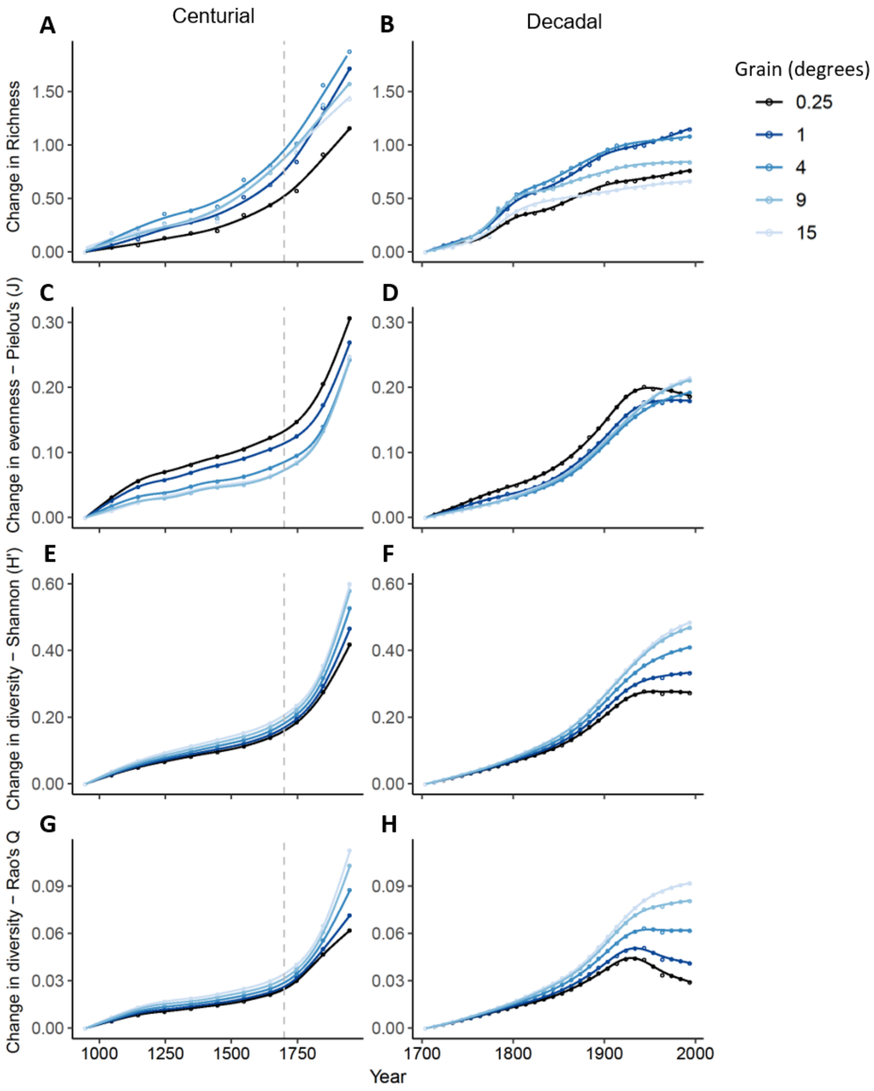


**Fig. S3**. **Spatial scaling of ecosystem diversity trends (*α* diversity).** Plots show the GAM-fitted trends for change in **(A-B)** within-cell ecosystem richness (change in mean numbers of ecosystem types per cell), **(C-D)** Pielou's evenness (J), **(E-F)** Shannon diversity Index (H') **(G-H)**, and Rao's quadratic entropy Index (RaoQ) for five different grains of analysis (0.25º, 1º, 4º, 9º and 15º). Each point on the graph represents the averaged change values across all cells of a given grain found over a 100yr and 10yr period (plotted in the mid-point of the century and decade) and are shown relative to the 10th century and to 1700, respectively. The smoothed lines represent fitted GAMs regressions.

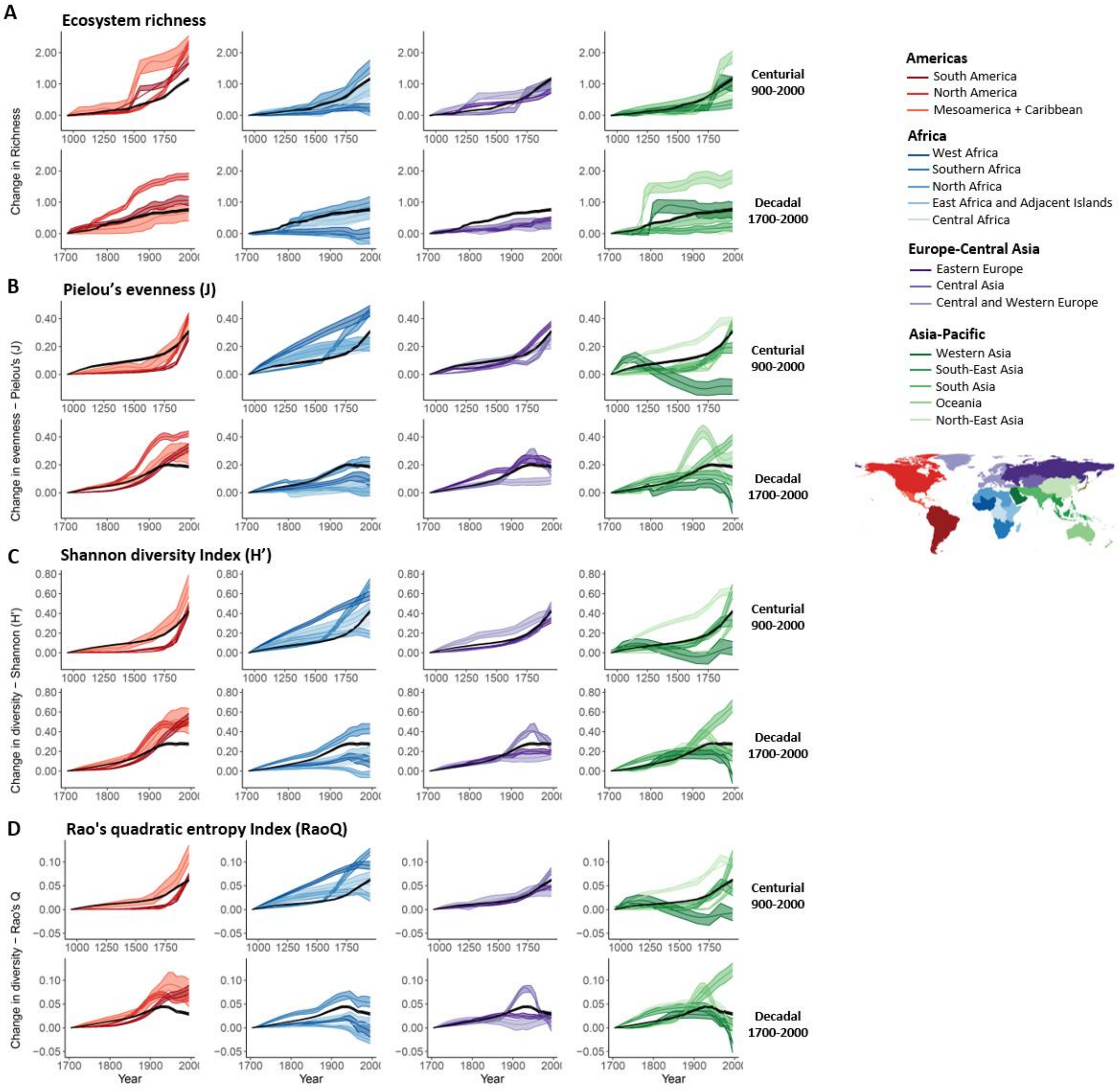


**Fig. S4. Ecosystem diversity trends ($\alpha$ diversity) for each IPBES sub-region.** Average regional changes of **(A)** within-sample ecosystem richness (change in mean numbers of ecosystem types per cell), **(B)** Pielou's evenness (J), **(C)** Shannon diversity Index (H'), and **(D)** Rao's quadratic entropy Index (RaoQ) across cells (0.25º x 0.25º) found over a 100yr and 10yr period, and shown relative to the 10$^{th}$ century and to 1700, respectively. All smoothed lines represent fitted GAM regressions; the shading represents the interdecile range calculated from 1000 random draws (each draw containing 2095 random sampling cells). Colours represent different regions; black lines show GAM-fitted global trends.

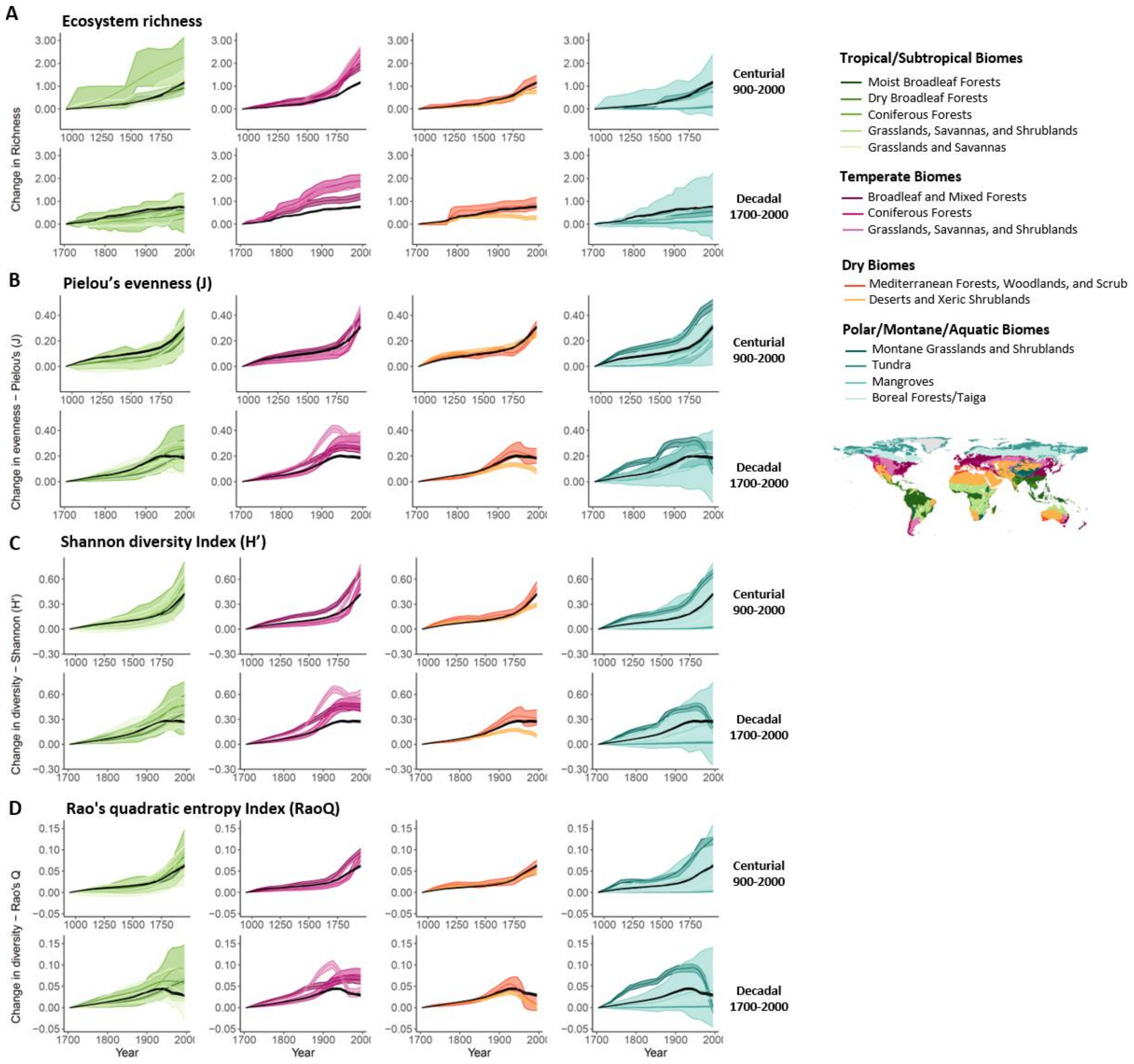


**Fig. S5.** Same as **Fig. S4,** but showing average change within each WWF Biome.

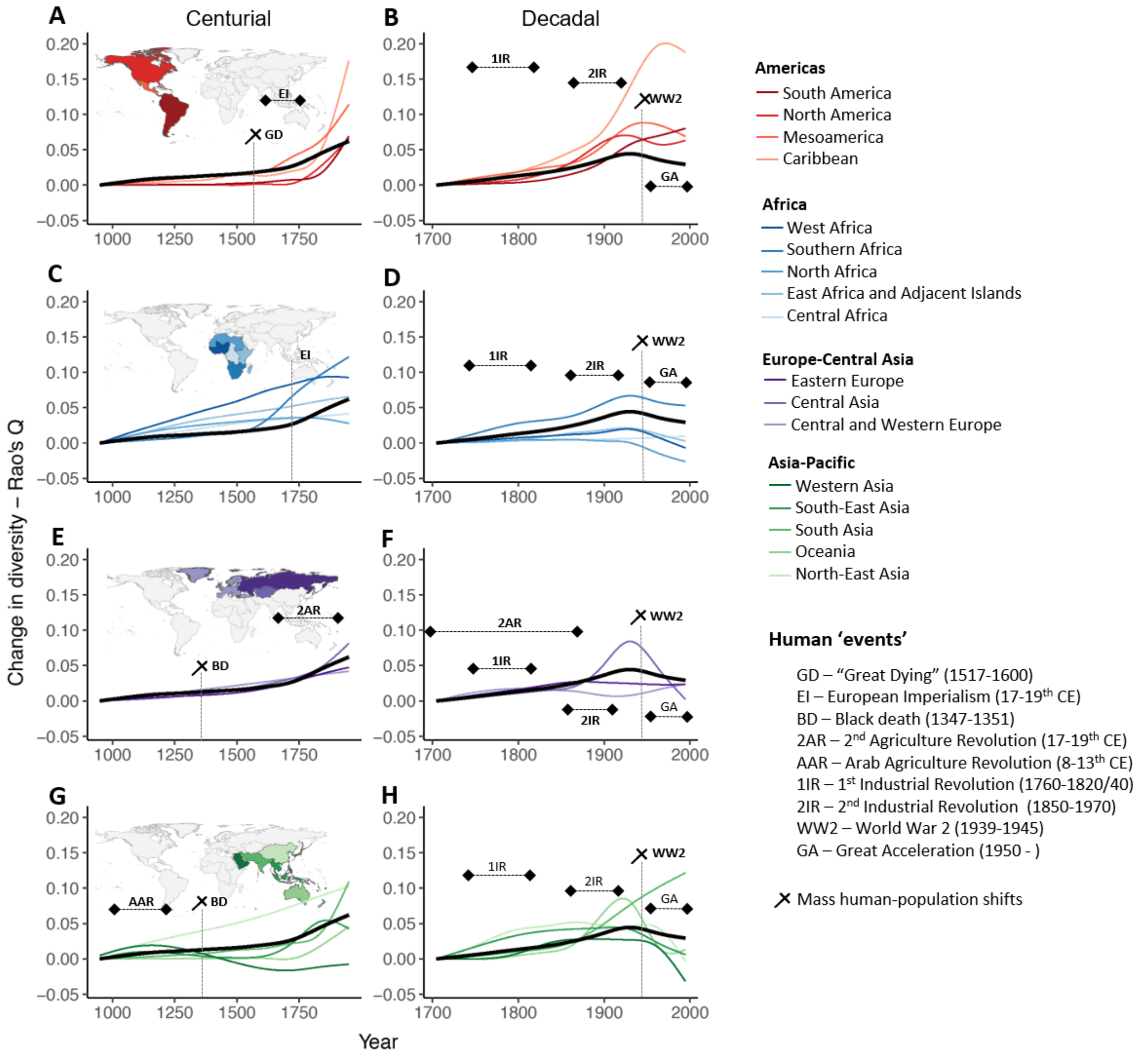


**Fig. S6. Coincidence between major socio-economic events and trajectories of local ecosystem diversity change (Rao's Q Index)** across **(A-B)** the Americas, **(C-D)** Africa, **(E-F)** Europe-Central Asia and **(G-H)** Asia-Pacific sub-regions. Each point on the graph represents the averaged values across all cells (0.25º x 0.25º) found over a 100yr and 10yr period (plotted in the mid-point of the century and decade) and shown relative to the 10th century and to 1700, respectively. The smoothed lines represent fitted GAMs regressions. Colours represent different regions; black lines show GAM-fitted global trends.

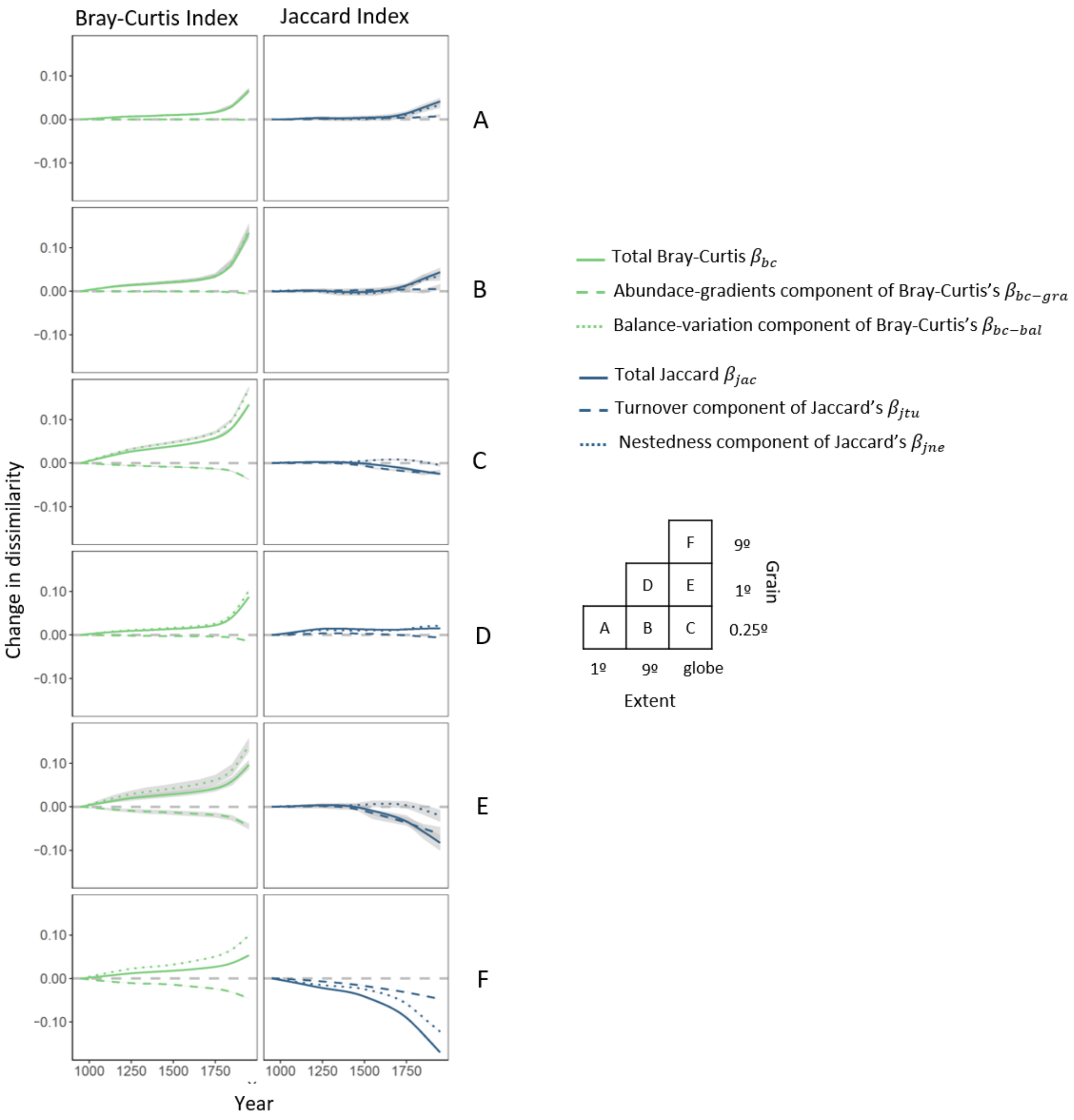


**Fig. S7. Decomposition of spatial $\beta$ diversity centurial patterns.** Plots show the different components of dissimilarity trends shown in Fig. 3. The Bray-Curtis Index ($\beta_{bc}$) is split into balanced variation ($\beta_{bc-bal}$) and abundance gradient ($\beta_{bc-gra}$) components, and the Jaccard Index ($\beta_{jac}$) into turnover ($\beta_{jtu}$) and nestedness ($\beta_{jne}$) components.

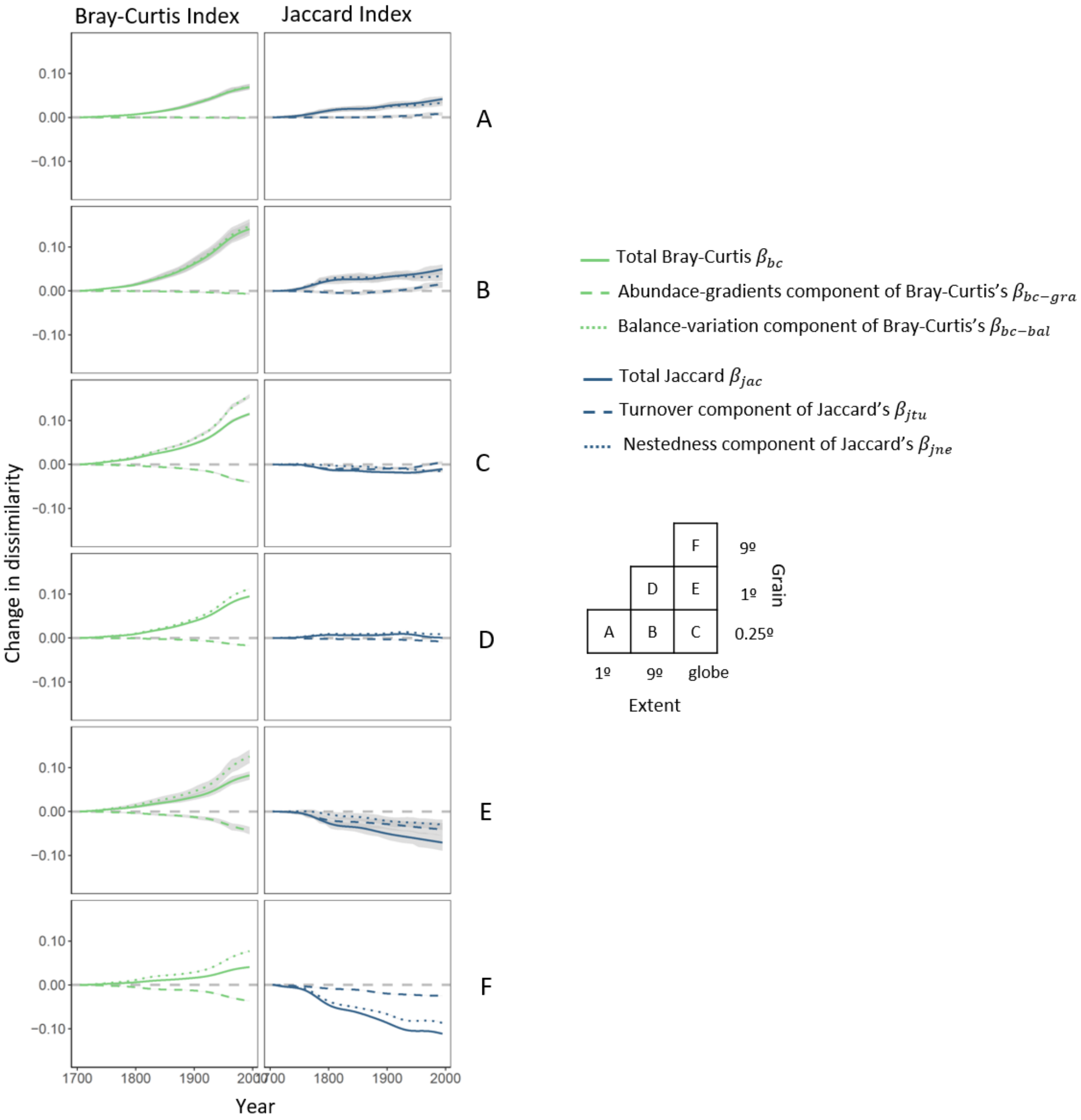


**Fig. S8.** Same as **Fig. S7,** but showing decadal trends.

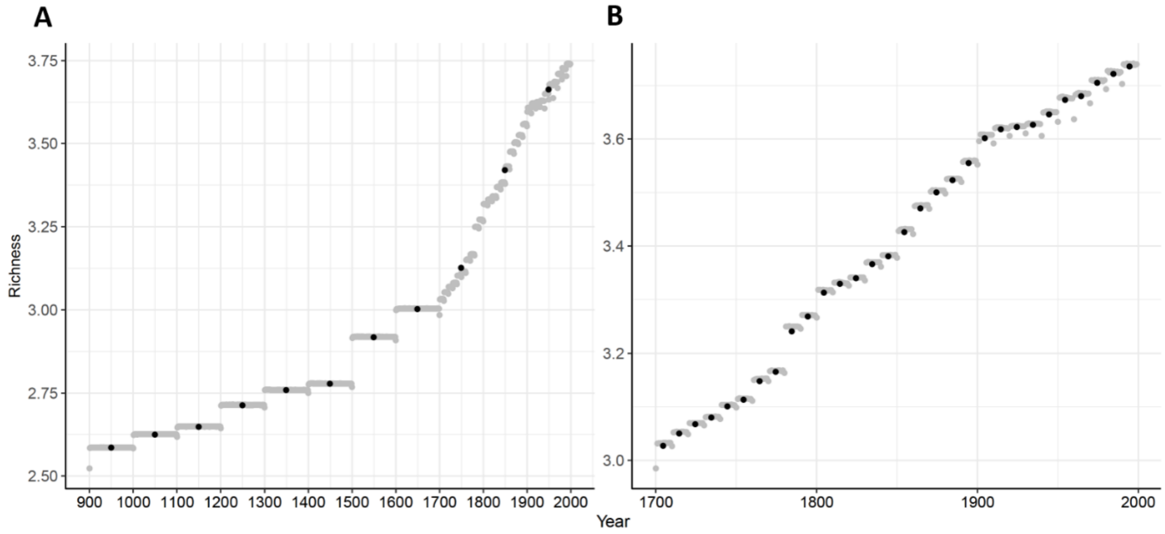


**Fig. S9. LUH2 input data temporal resolution**. The original land-use data behind the LUH2 model had an underlying temporal resolution of every 100 years from 900 to 1700 and every 10 years from 1700 to 2000. An algorithm was used by the LUH2 developers to transform century- and decadal-long estimates to annual estimates based on a set of assumptions (*6*). In our analysis, we re-aggregated to **(A)** centurial and **(B)** decadal temporal resolutions (e.g., for ecosystem richness, shown here; black points), because individual year values (grey points) are not independent of one another. For centurial analyses, we also calculated mean values for 100-year intervals after 1700.

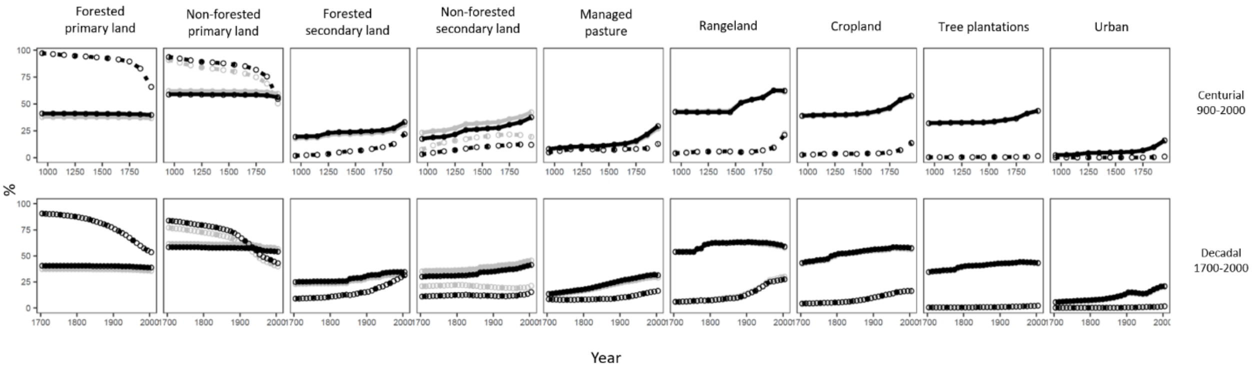


**Fig. S10. The impact of North Africa and Western Asia cells on global estimates of ecosystem coverage**. Plots shows average presence (i.e. % of cells where each ecosystem type is present - solid line) and coverage (i.e. % of the cell occupied by the ecosystem dashed line) of ecosystems across all cells (grey line) and when excluding North Africa and Western Asia cells (black line). All lines represent average values across 0.25º x 0.25º grid cells. Where grey lines cannot be seen, they lie beneath the black lines, indicating minimal effect of North African and Western Asian cells. The only notable difference relates to the balance of primary and secondary non-forested land, which is a true reflection of the predominance of dryland ecosystems in this region.

**Data S1. Average change in the presence and coverage of the different ecosystem types from 900-2000.** Column "*version*" refers to change across (V1) all cells or (V2) excluding North Africa and Western Asia cells.

**Data S2. Global and regional (IPBES sub-regions and WWF Biomes) changes in local ecosystem diversity ($\alpha$ diversity).** Table includes all data shown in Fig. 1 and 2.

**Data S3. Temporal trends in spatial $\beta$ diversity of ecosystems.** Column "*n_pairs*" refers to the total (when "*n_sim*" =1) or average across draws (when "*n_sim*" =100) number of pairwise comparisons for each combination of grain and extent. Table includes all data shown in Fig. 3.
